# Whole blood transcriptome profile identifies motor neurone disease (MND) RNA biomarker signatures

**DOI:** 10.1101/2024.09.27.615372

**Authors:** Sulev Kõks, Karin Rallmann, Mari Muldmaa, Jack Price, Abigail L Pfaff, Pille Taba

## Abstract

Blood-based biomarkers for motor neuron disease are needed for better diagnosis, progression prediction, and clinical trial monitoring. We used whole blood-derived total RNA and performed whole transcriptome analysis to compare the gene expression profiles in (motor neurone disease) MND patients to the control subjects. We compared 42 MND patients to 42 aged and sex-matched healthy controls and described the whole transcriptome profile characteristic for MND. In addition to the formal differential analysis, we performed functional annotation of the genomics data and identified the molecular pathways that are differentially regulated in MND patients. We identified 12,972 genes differentially expressed in the blood of MND patients compared to age and sex-matched controls. Functional genomic annotation identified activation of the pathways related to neurodegeneration, RNA transcription, RNA splicing and extracellular matrix reorganisation. Blood-based whole transcriptomic analysis can reliably differentiate MND patients from controls and can provide useful information for the clinical management of the disease and clinical trials.

## 1. Introduction

Motor neurone disease (MND) is a group of chronic sporadic and familial disorders characterised by progressive degeneration of motor neurons [1]. The disease is caused by the degeneration of the upper, lower, or both motor neurones. The prognosis of MND depends upon the age at onset and the area of the central nervous system affected [2]. Based on the site of origin and the severity of neurological involvement, four main subtypes of MND have been described: amyotrophic lateral sclerosis (ALS), progressive bulbar palsy (PBP), progressive muscular atrophy (PMA), and primary lateral sclerosis (PLS) [3].

ALS is the most common form of MND. ALS and MND are commonly used interchangeably or as synonyms. ALS is also known as Lou Gehrig’s disease or Charcot disease [1]. ALS is an adult-onset, progressive, neurodegenerative disorder involving the large motor neurons of the brain and the spinal cord. It produces a characteristic clinical picture with weakness and wasting of the limbs and bulbar muscles, leading to death from respiratory failure within five years.

The degeneration of motor neurons is irreversible, and apparently, it starts many years before the clinical features emerge. Therefore, reliable biomarkers from easily accessible tissues are needed for earlier diagnosis and better prediction of the progression of the disease. The molecular pathology underlying MND relies on genetic variants described in at least 100 different genes to date and on the overlay of the transcriptomic changes [4, 5]. The pathogenesis of the disease involves oxidative stress, inflammation, ER stress with protein aggregation, autophagy and aberrant RNA processing [5, 6]. Familial and sporadic forms of MND can be distinguished based on the evidence of genetic variants and family history [7]. However, only about 20% of MND cases can be explained by known genetic variations [8]. In addition to the well-known genes and their variants, we recently described an unexpectedly large number of exonisation of SINE-VNTR-Alu repeats (SVAs) in the motor cortex [9]. SVAs are known to alter splicing, and several of these elements have been associated with disease through such mechanisms [10, 11]. This indicates the significant role that noncoding or dark genomes can play in the pathogenesis of complex diseases. Moreover, analysis of the whole transcriptome gives an excellent functional opportunity to explore the molecular changes at different stages of diseases, making it a suitable tool for biomarkers [10]. Indeed, transcriptomic analysis can be performed from any biological material, like blood or cerebrospinal fluid and can be used for different conditions [5, 12, 13]. Transcriptomic analysis helps to understand the effect of DNA variants, especially for the splicing-altering variants.

Post-mortem tissue analysis for chronic diseases is always an option to identify molecular patterns in the affected tissues, and this can help to classify the different pathogenic mechanisms [6]. However, using peripheral tissues, like blood, skin, or saliva, allows molecular profiling during the disease’s progression and real-life monitoring of pathogenic changes [12, 14, 15]. In the case of MND, several previous studies have been performed to analyse the transcriptomic profile of the blood [16, 17]. In one example, whole blood-derived RNA (PAXgene tubes) was used for microarray analysis; in another, PBMC-derived RNA was used for RNA sequencing. The present study aimed to perform whole transcriptome analysis from the whole-blood (Tempus tubes) derived RNA and to identify the whole blood transcriptomic profile by comparing MND patients to the age and sex-matched healthy controls.

## 2. Materials and Methods

The whole blood was collected from 42 MND patients and 42 healthy controls using Tempus Blood RNA collection tubes (Thermo Fisher Scientific). Neurologists recruited MND patients, and the subtype of the MND was confirmed. Healthy controls were recruited among the visitors referred to the blood analysis who did not have chronic diseases. The control samples were ideal controls without any neurological condition or major chronic illness and were age- and sex-matched to the MND group (complete information is given in Supplementary Table 1).

The RNA was isolated from whole blood using Tempus Spin Isolation Kit (Thermo Fisher Scientific). After initial quality control and quantification, RNA was used for the total RNA sequencing necessary for the whole transcriptome analysis.

Total RNA sequencing was performed in all 84 samples at the Genomics Core Facility at Murdoch University, Perth, WA. Illumina paired-end 2×100bp read length using NovaSeq 6000. The NovaSeq Control Software v1.7.5 and Real-Time Analysis (RTA) v3.4.4 performed real-time image analysis. RTA performs real-time base calling on the NovaSeq instrument computer. The Illumina DRAGEN BCL Convert 07.021.624.3.10.8 pipeline generated the sequence data. The FASTQ files were analysed using salmon 1.10.3 by using the reference genome GRCh38 [18]. Salmon counts were imported to the R studio using the *tximeta* package [19]. Differential whole transcriptome analysis was performed with the *DESeq2* package [20]. No fold-change filtering was applied, but the False Discovery Rate (FDR) was set at 0.05 to adjust for multiple testing. The functional annotation of the differential gene expression was performed with the packages *ReactomePA*, *clusterProfiler* and *DOSE* [21–23].

To perform a pair-wise analysis of individual genes between MND and healthy controls, we applied the two-tailed Wilcoxon rank-sum test implemented in the function compare_means() of the package *ggpubr* [24]. We generated a list of all known MND genes using the OMIM catalogue and identified 97 genes that are directly connected to the MND or its subtypes. This list extracted normalised counts from the salmon quant files and made boxplots with pairwise comparisons. Plots were generated using *ggplot2* version 3.5.1 and *ggpubr* version 0.6.0 packages. Statistical analysis was performed with R software version 4.4.0 (https://www.R-project.org) and RStudio Version 2023.06.0+421.

## 3. Results

### 3.1. Description of the study cohort

Between 2013 and 2018, a total of 84 patients (42 MND patients and 42 healthy control patients without any chronic diseases) were enrolled in the study and signed written informed consent. Inclusion criteria for MND patients were the diagnosis of probable or definitive MND based on El Escorial Criteria and the absence of a positive family history. For the healthy controls, we excluded patients with any chronic diseases, especially any neurologic, rheumatological, haematological, or oncological conditions. In addition, treatment with biologics or chemotherapy was also excluded. The blood samples were collected into Tempus Blood RNA tubes and stored according to the manufacturer’s instructions. The research was conducted with the approval of the University of Tartu Research Ethics Committee, and all participants provided written informed consent. The comprehensive patient selection process leveraged hospital records, neurologist consultations, and the Estonian Health Insurance Fund’s national health data repository. The general characteristics of the population are reported in Table 1. The median age was 65.6 (standard deviation 9.3) years, and most subjects were female (69%). No patient reported a positive family history of MND; therefore, all the participants had sporadic forms, and all patients received standard MND therapy with riluzole. The most frequent clinical subtype was the classic ALS (86%). Spinal symptoms were present the most commonly (60%).

**Table 1.**
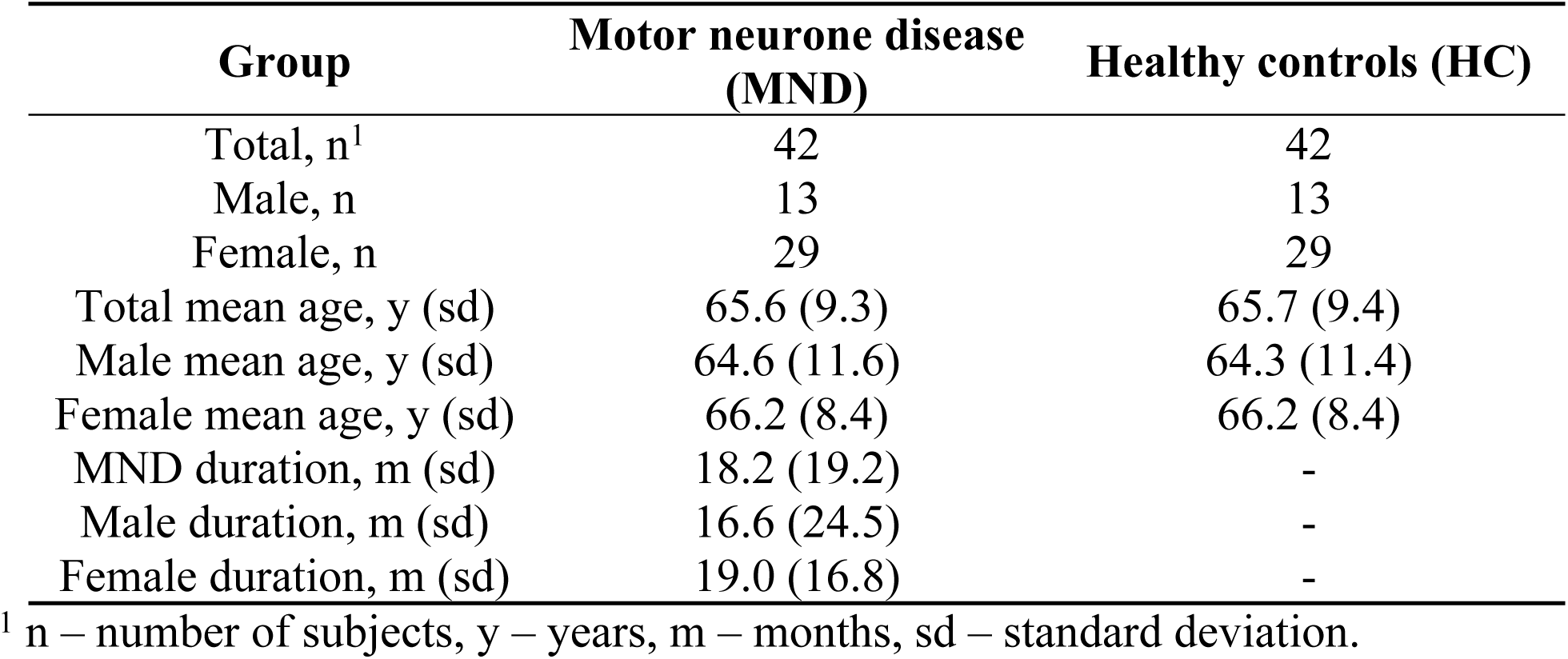
General characteristics of the study cohort.

| **Group** | **Motor neurone disease**<br>**(MND)** | **Healthy controls (HC)** |
| --- | --- | --- |
| Total, n<sup>1</sup> | 42 | 42 |
| Male, n | 13 | 13 |
| Female, n | 29 | 29 |
| Total mean age, y (sd) | 65.6 (9.3) | 65.7 (9.4) |
| Male mean age, y (sd) | 64.6 (11.6) | 64.3 (11.4) |
| Female mean age, y (sd) | 66.2 (8.4) | 66.2 (8.4) |
| MND duration, m (sd) | 18.2 (19.2) | - |
| Male duration, m (sd) | 16.6 (24.5) | - |
| Female duration, m (sd) | 19.0 (16.8) | - |

<sup>1</sup> n – number of subjects, y – years, m – months, sd – standard deviation.

### 3.2. Whole blood RNA sequencing

RNA sequencing resulted in at least 50 million paired reads per sample. Salmon was used to quantify transcript abundances from fastq files. *Tximeta* was used to import the resulting quant files, and gene-level summarisation was used for the *DESeq2* workflow. Healthy controls were compared to the MND RNA-seq results, and we identified 12,972 genes differentially expressed (FDR < 0.05) in the blood of MND patients. The top 30 differentially regulated genes are shown in Table 2. Out of these 12,972 genes, 8,008 were upregulated, and 4,964 were down-regulated (Supplementary Table 2). A heat map with all 12,972 genes is shown as supplementary in Figure 1, and it shows a clear separation of MND patients from the healthy controls. A smaller heatmap with the top 100 genes is shown in Figure 1, and a volcano plot is shown in Figure 2. The heatmap with 100 genes shows a consistent and clear separation of the MND from the healthy controls. This remarkable finding shows that a disease highly specific to the central nervous system can be differentiated from controls by the blood transcriptome profile.

**Table 2.** Differentially expressed genes in the blood of MND patients compared to healthy controls. The top 30 genes are shown sorted by the FDR-adjusted p-value.

| Ensembl ID | logFC | p-adj | Gene name | Gene symbol |
| --- | --- | --- | --- | --- |
| ENSG00000202354 | 6.49 | 1.25E-136 | RNA, Ro60-associated Y3 | RNY3 |
| ENSG00000201098 | 7.26 | 4.63E-119 | RNA, Ro60-associated Y1 | RNY1 |
| ENSG00000282885 | 3.29 | 3.44E-111 | novel transcript | lnc-NEMF-1 |
| ENSG00000091986 | -6.81 | 4.27E-101 | coiled-coil domain containing 80 | CCDC80 |
| ENSG00000011465 | -7.48 | 9.56E-99 | decorin | DCN |
| ENSG00000118523 | -4.88 | 5.04E-97 | cellular communication network factor 2 | CCN2 |
| ENSG00000164692 | -7.26 | 1.58E-95 | collagen type I alpha 2 chain | COL1A2 |
| ENSG00000108821 | -6.43 | 1.04E-93 | collagen type I alpha 1 chain | COL1A1 |
| ENSG00000128591 | -5.83 | 8.20E-93 | filamin C | FLNC |
| ENSG00000138131 | -5.55 | 1.70E-87 | lysyl oxidase like 4 | LOXL4 |
| ENSG00000113739 | -7.73 | 2.01E-86 | stanniocalcin 2 | STC2 |
| ENSG00000199568 | 9.05 | 1.07E-80 | RNA, U5A small nuclear 1 | RNU5A-1 |
| ENSG00000248527 | 3.20 | 6.12E-80 | MT-ATP6 pseudogene 1 | MTATP6P1 |
| ENSG00000087245 | -6.83 | 6.87E-80 | matrix metalloproteinase 2 | MMP2 |
| ENSG00000186340 | -7.55 | 2.94E-79 | thrombospondin 2 | THBS2 |
| ENSG00000115414 | -5.71 | 1.45E-78 | fibronectin 1 | FN1 |
| ENSG00000150459 | -0.79 | 3.01E-77 | Sin3A associated protein 18 | SAP18 |
| ENSG00000212283 | 5.02 | 4.23E-77 | small nucleolar RNA, C/D box 89 | SNORD89 |
| ENSG00000111799 | -6.88 | 1.03E-75 | collagen type XII alpha 1 chain | COL12A1 |
| ENSG00000199631 | 7.52 | 2.52E-73 | small nucleolar RNA, C/D box 33 | SNORD33 |
| ENSG00000144810 | -6.44 | 3.84E-73 | collagen type VIII alpha 1 chain | COL8A1 |
| ENSG00000164761 | -6.80 | 1.42E-72 | TNF receptor superfamily member 11b | TNFRSF11B |
| ENSG00000115963 | -6.65 | 2.38E-70 | Rho family GTPase 3 | RND3 |
| ENSG00000115461 | -7.69 | 9.88E-70 | insulin like growth factor binding protein 5 | IGFBP5 |
| ENSG00000126214 | 0.95 | 1.05E-63 | kinesin light chain 1 | KLC1 |
| ENSG00000186660 | -0.54 | 1.50E-63 | ZFP91 zinc finger protein, E3 ubiquitin ligase | ZFP91 |
| ENSG00000142156 | -2.90 | 1.73E-62 | collagen type VI alpha 1 chain | COL6A1 |
| ENSG00000238961 | 5.94 | 3.17E-62 | small nucleolar RNA, H/ACA box 47 | SNORA47 |
| ENSG00000166923 | -7.07 | 2.09E-61 | gremlin 1, DAN family BMP antagonist | GREM1 |
| ENSG00000281501 | 1.98 | 5.08E-61 | SEPSECS antisense RNA 1 | SEPSECS-AS1 |

**Figure 1.**
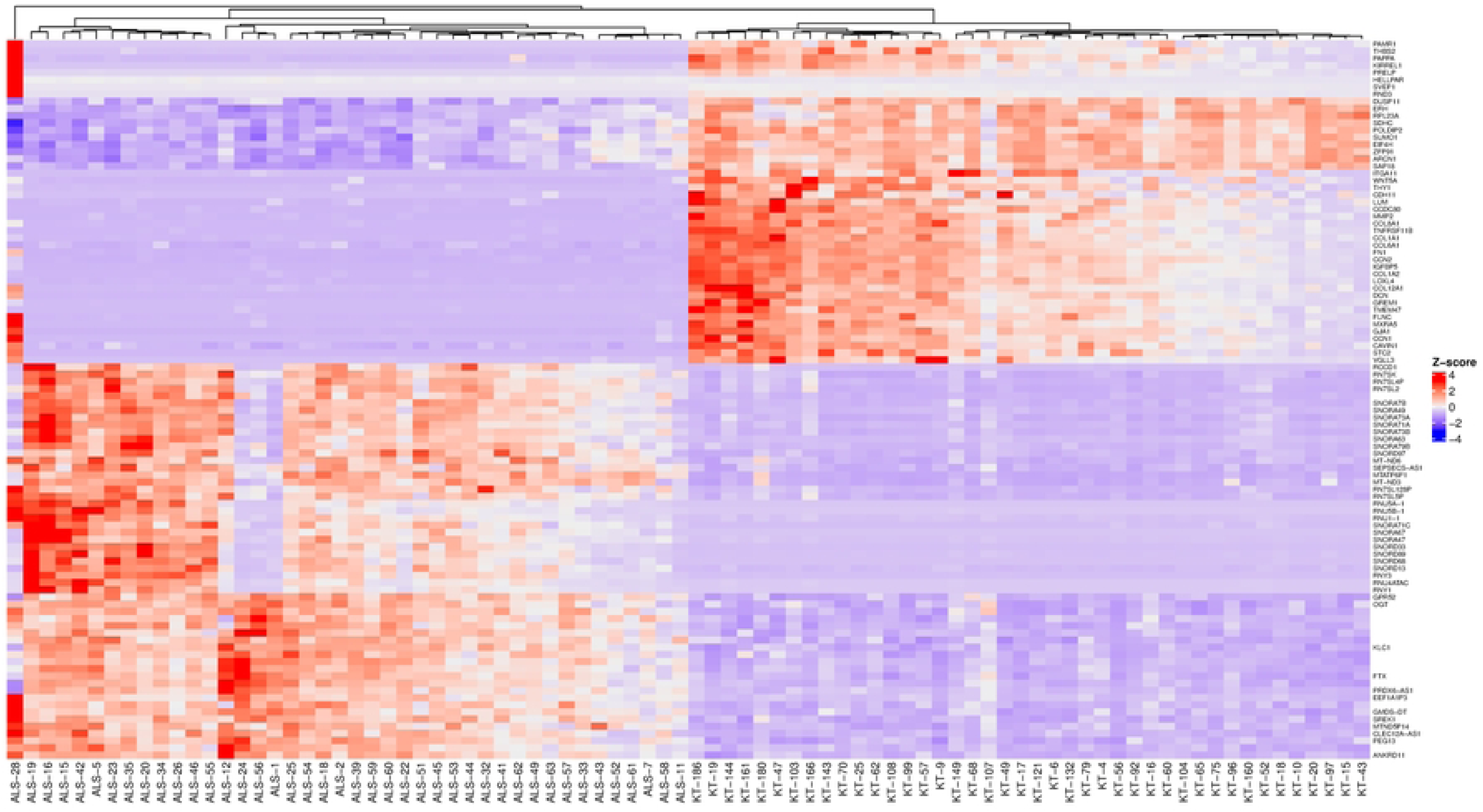
Heatmap of the 100 differentially expressed genes with the highest statistical significance. Samples with “ALS” designate the MND group, and “KT” designate healthy controls.

**Figure 2.**
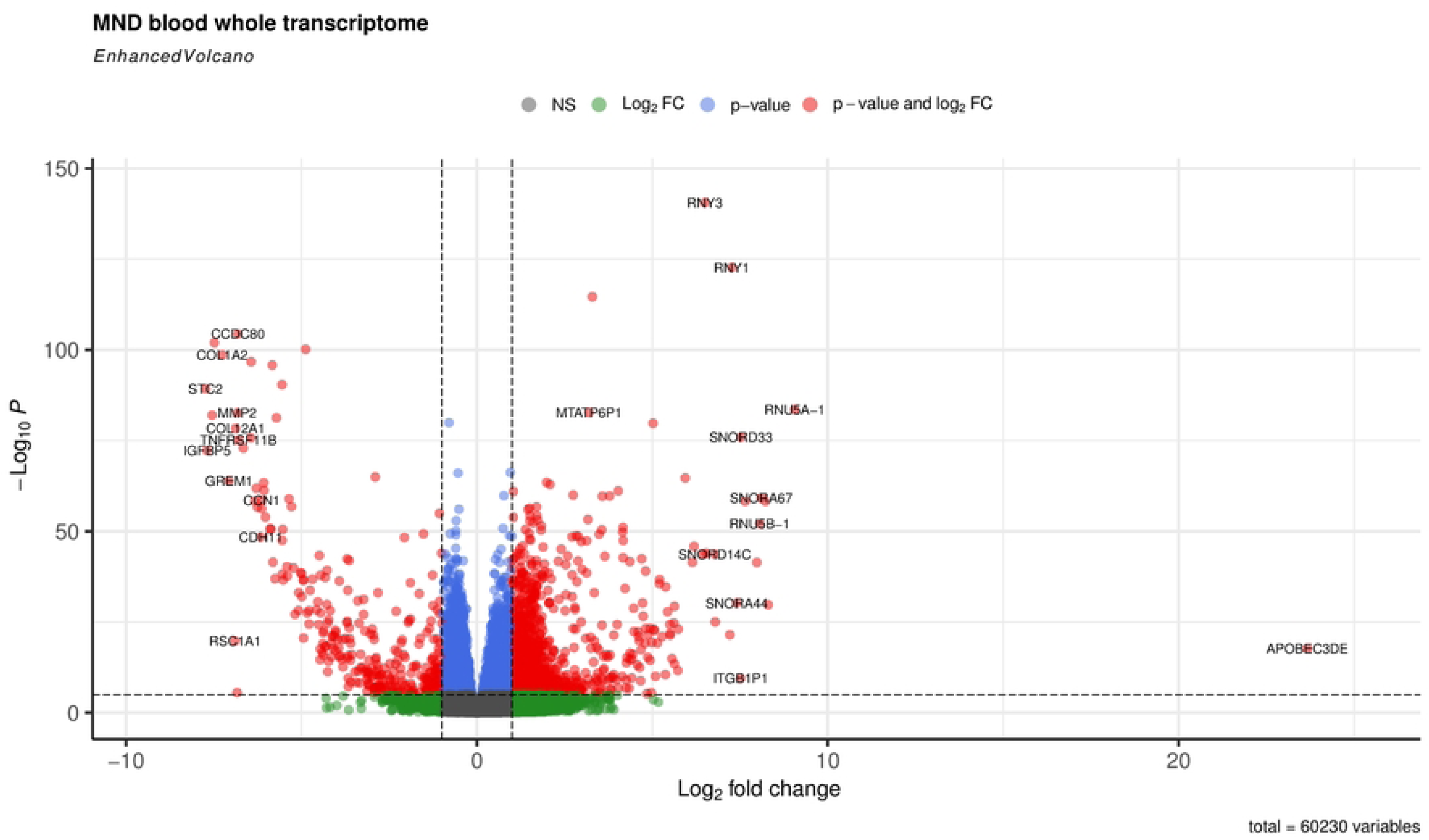
Volcano plot of the whole transcriptome data from the blood on controls and MND patients The de-fault cut-off for log2FC is >|2|, and the default for P-value is 10e-6. Dashed lines represent these values. Red dots represent genes meeting both cut-off criteria; green dots meet only the log2FC cut-off, and blue dots indicate genes meeting only the P-value cut-off.

### 3.3. Pairwise analysis of known MND genes

In addition to the whole transcriptome analysis, we performed a pairwise (MND versus healthy controls) study of 97 known MND genes (a list of the genes is provided in Supplementary Table 3) and 30 top-regulated genes from the DESeq2 analysis. All results are shown in Supplementary Figure 2, and partial results are in Figures 3 and 4. Interestingly, some MND-related genes are upregulated (ALS2, NEK1, ATXN2), while others are downregulated (SOD1, UBQLN2 aka ALS15) in patients. In addition, FUS and ANXA11 were upregulated, and ANG was downregulated in patients (Figure 4 A, B, C, D, E, F). Moreover, the DESeq2 top genes RNY3, RNY1, and ENSG0000282885 were highly upregulated in patients with almost no expression in control subjects (Figure 4 G, H, I). At the same time, other DESeq2 top genes, CCDC80, DCN, and CCN2, were highly expressed in controls, and their expression was almost missing in patients’ blood (Figure 4 J, K, L). These examples indicate that there are many high fold-change difference genes with almost no expression in one group and very high expression in another, and these genes have very high potential to be a transcriptional biomarker for the MND.

**Figure 3.**
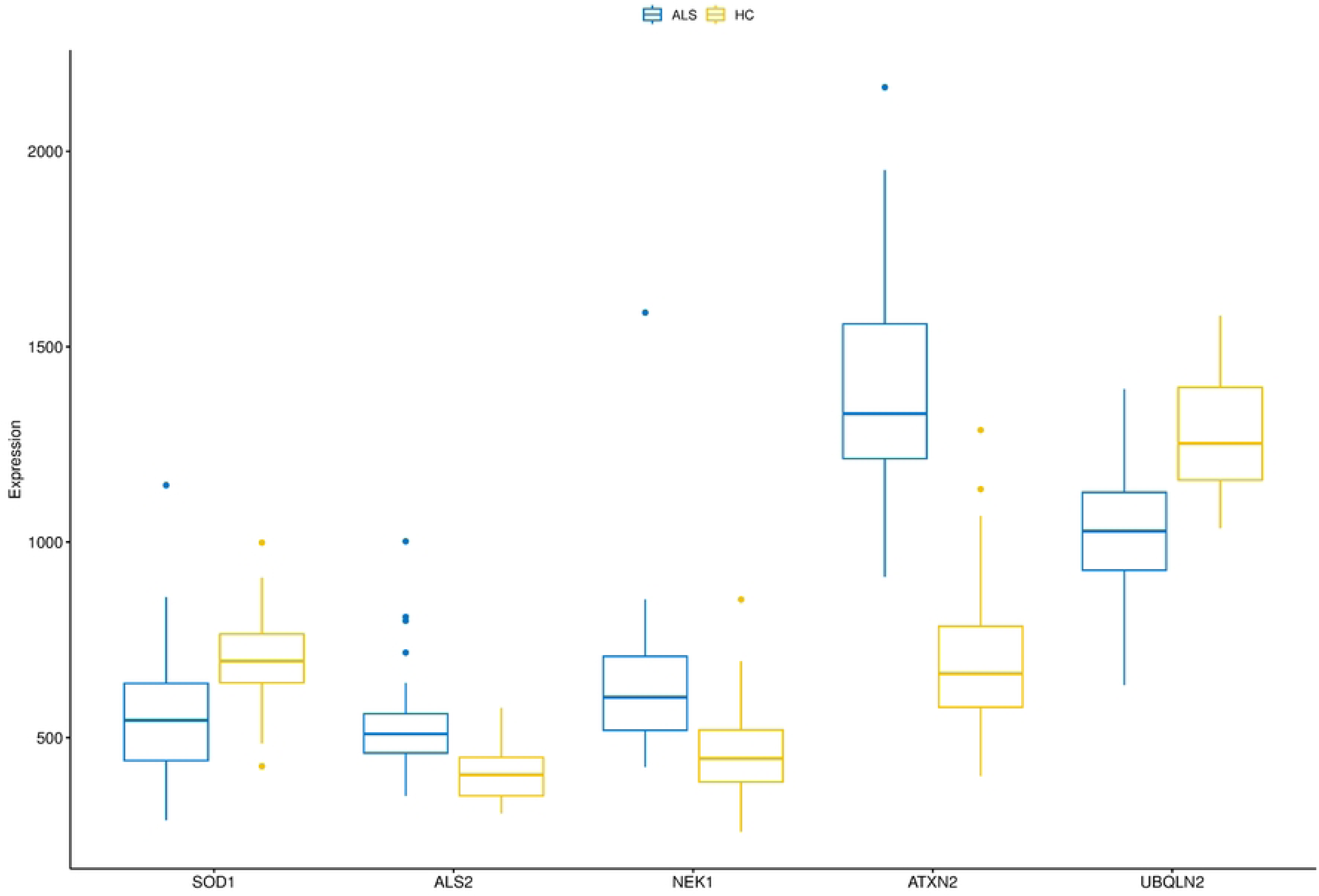
A combined boxplot of five MND-related genes and their expression levels in the blood of MND patients and controls gives comparative blood expression levels for these selected genes. Pairwise statistical comparisons are shown in Figure 4 and in supplementary Figure 2.

**Figure 4.**
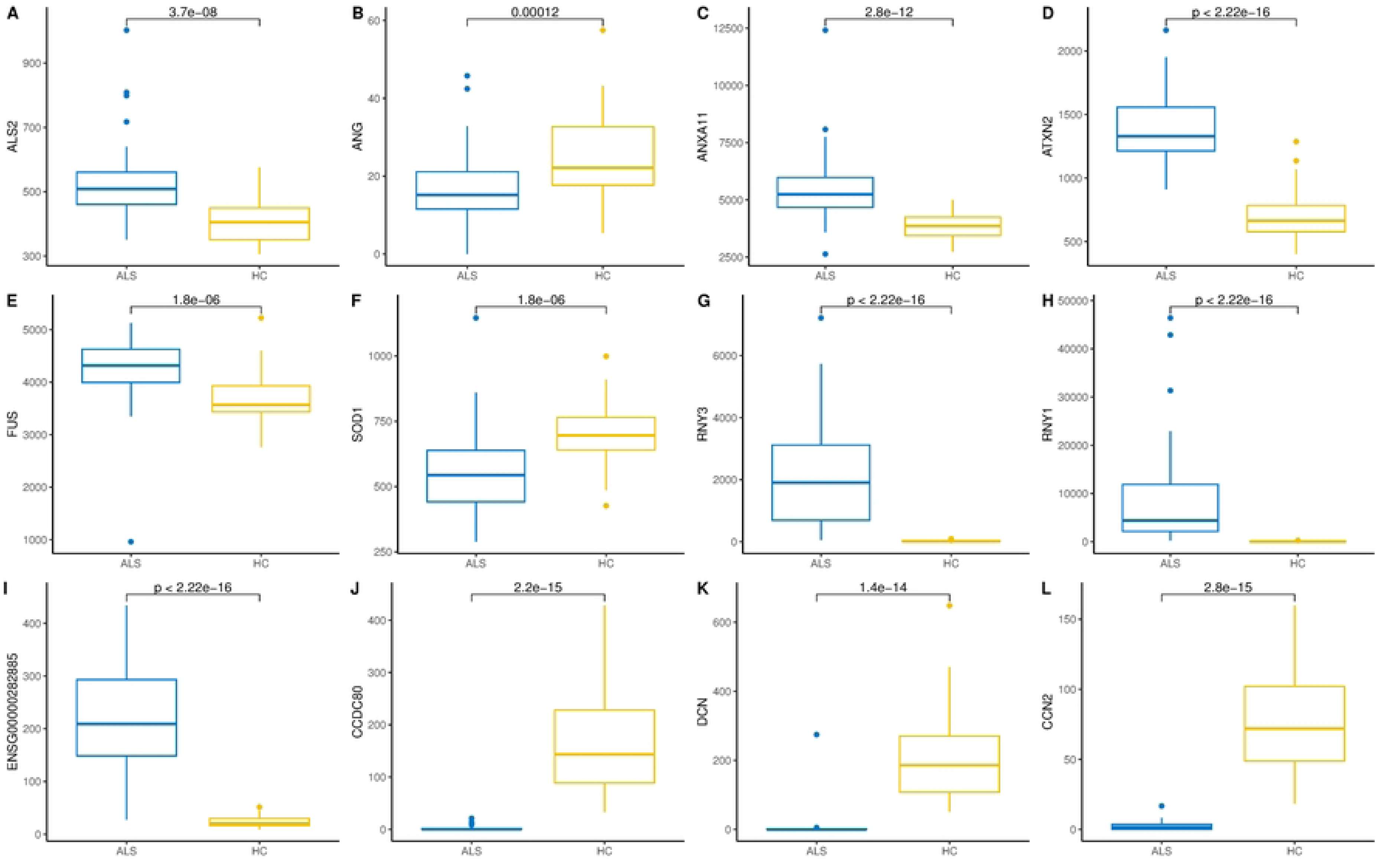
Pairwise comparison (Wilcoxon rank-sum test) and boxplots of six MND-related genes (A-F) and six of the most significant differentially expressed genes (G-L) in the blood of MND pa-tients and controls. The Y-axis shows gene expression in normalised counts.

The pairwise analysis of all 97 MND genes indicated that some well-known MND genes weren’t differentially expressed in the blood (boxplots are in Supplementary Figure 2). Out of all 97 genes, 38 (39%) of them AMFR, AR, ATX3, BICD2, C9orf72, CHRNA3, DAO, DCTN1, DNAJC7, ERBB4, HNRNPA2B1, IGFALS, KIF5A, LGALSL, LRP12, MAPT, MOBP, NEFH, OPTN, PAH, PON1, PON2, PON3, PRPH, PSEN1, SARM1, SCYL1, SETX, SLC1A2, SLC52A3, SMN1, SMN2, SQSTM1, TARDBP, TRPM7, TUBA4A, VRK1, VSX1, were not differentially expressed between patients and controls. Fourteen genes of these 38 genes were not expressed in blood. Most of these genes that were not differentially expressed had excellent expression levels in the blood. AMFR has an expression level of 1,800 normalised counts, C9orf72 has 1,500 normalised counts, PSEN1 has an expression at 2,500 normalised counts, TARDBP has an average gene expression of 1,600 normalised counts, SQSTM1 has an expression level of 3,100 normalised counts. Therefore, all these genes are highly expressed in the whole blood, but their expression level is not dependent on the disease status.

### 3.4. Functional annotation of differentially expressed genes

Functional annotation of differentially expressed genes indicated statistically significant activation of several human disease pathways (Table 3, full version provided in Supplementary Table 4). Remarkably, three neurodegenerative diseases were at the top of the table of the KEGG pathways: Parkinson’s disease, prion disease, and amyotrophic lateral sclerosis (Figure 5). In addition, several pathways involved in the pathogenesis of neurodegeneration were also activated. These included protein processing in the endoplasmic reticulum, proteasome, lysosome and ubiquitin-mediated proteolysis.

**Table 3.** KEGG pathways that are enriched in the blood transcriptome of MND patients.

| Category | Subcategory | ID | Description | Gene Ratio | Bg Ratio | P-adjusted |
| --- | --- | --- | --- | --- | --- | --- |
| Human Diseases | Neurodegenerative disease | hsa05012 | Parkinson disease | 165/3781 | 271/8843 | 2.36E-07 |
| Human Diseases | Cardiovascular disease | hsa05415 | Diabetic cardiomyopathy | 130/3781 | 205/8843 | 2.36E-07 |
| Human Diseases | Neurodegenerative disease | hsa05020 | Prion disease | 167/3781 | 278/8843 | 3.14E-07 |
| Human Diseases | Neurodegenerative disease | hsa05014 | Amyotrophic lateral sclerosis | 212/3781 | 371/8843 | 5.97E-07 |
| Metabolism | Energy metabolism | hsa00190 | Oxidative phosphorylation | 92/3781 | 138/8843 | 5.97E-07 |
| Organismal Systems | Environmental adaptation | hsa04714 | Thermogenesis | 143/3781 | 235/8843 | 5.97E-07 |
| Genetic Information Processing | Folding, sorting and degradation | hsa04141 | Protein processing in endoplasmic reticulum | 109/3781 | 170/8843 | 5.97E-07 |
| Human Diseases | Neurodegenerative disease | hsa05022 | Pathways of neurodegeneration - multiple diseases | 265/3781 | 483/8843 | 1.07E-06 |
| Human Diseases | Cancer: overview | hsa05208 | Chemical carcinogenesis - reactive oxygen species | 136/3781 | 226/8843 | 2.72E-06 |
| Human Diseases | Neurodegenerative disease | hsa05010 | Alzheimer disease | 217/3781 | 391/8843 | 5.09E-06 |
| Human Diseases | Neurodegenerative disease | hsa05016 | Huntington disease | 177/3781 | 311/8843 | 7.00E-06 |
| Genetic Information Processing | Translation | hsa03013 | Nucleocytoplasmic transport | 72/3781 | 108/8843 | 1.14E-05 |
| Genetic Information Processing | Folding, sorting and degradation | hsa03050 | Proteasome | 36/3781 | 46/8843 | 2.46E-05 |
| Cellular Processes | Transport and catabolism | hsa04142 | Lysosome | 82/3781 | 132/8843 | 0.0001 |
| Genetic Information Processing | Folding, sorting and degradation | hsa04120 | Ubiquitin mediated proteolysis | 87/3781 | 142/8843 | 0.0001 |
| Genetic Information Processing | Translation | hsa03010 | Ribosome | 102/3781 | 172/8843 | 0.0002 |
| Genetic Information Processing | Chromosome | hsa03082 | ATP-dependent chromatin remodeling | 73/3781 | 117/8843 | 0.0003 |
| Cellular Processes | Cell growth and death | hsa04110 | Cell cycle | 94/3781 | 158/8843 | 0.0003 |
| Genetic Information Processing | Replication and repair | hsa03030 | DNA replication | 28/3781 | 36/8843 | 0.0004 |
| Human Diseases | Infectious disease: bacterial | hsa05132 | Salmonella infection | 138/3781 | 251/8843 | 0.0009 |
| Metabolism | Glycan biosynthesis and metabolism | hsa00510 | N-Glycan biosynthesis | 37/3781 | 53/8843 | 0.0010 |
| Human Disease | Neurodegenerative disease | hsa05017 | Spinocerebellar ataxia | 84/3781 | 144/8843 | 0.0016 |

\* Bg – background.

**Figure 5.**
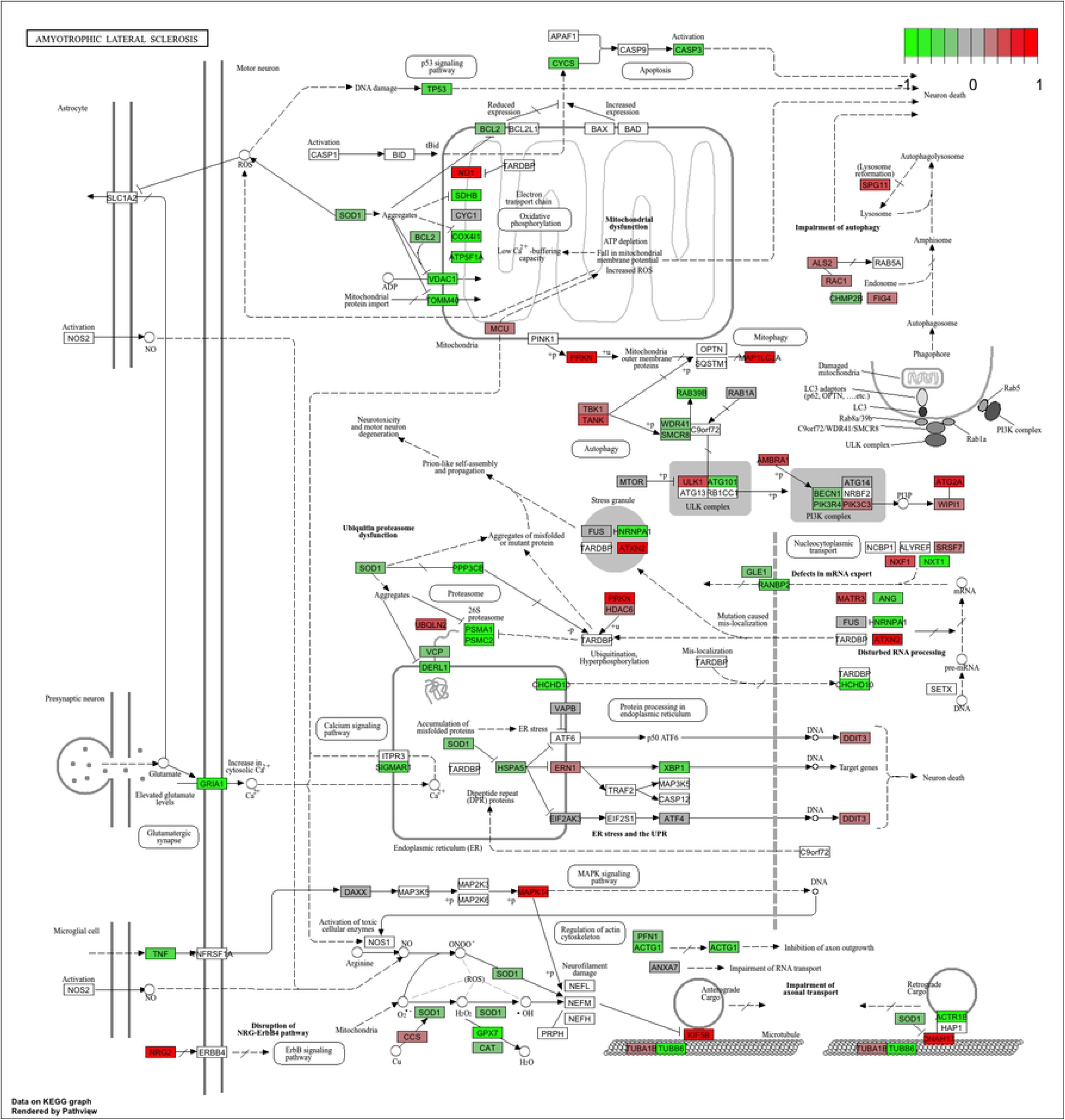
KEGG pathway “Amyotrophic Lateral Sclerosis” with the blood RNA gene expression data. Genes in green are downregulated, and genes in red are upregulated.

Reactome and GSEA analyses use more canonical pathways (Supplementary Tables 5 and 6). Reactome identified statistically significant enrichment of the mRNA splicing and transcription-related pathways in combination with cellular energetics pathways (mitochondria and respiratory electron transport) to be affected (Figure 6). GSEA analysis (Figure 7) identified statistically significant enrichment of sensory perception, olfactory signalling and many pathways related to the extracellular matrix reorganisation (collagen degradation, elastic fibre formation, assembly of collagen fibres).

**Figure 6.**
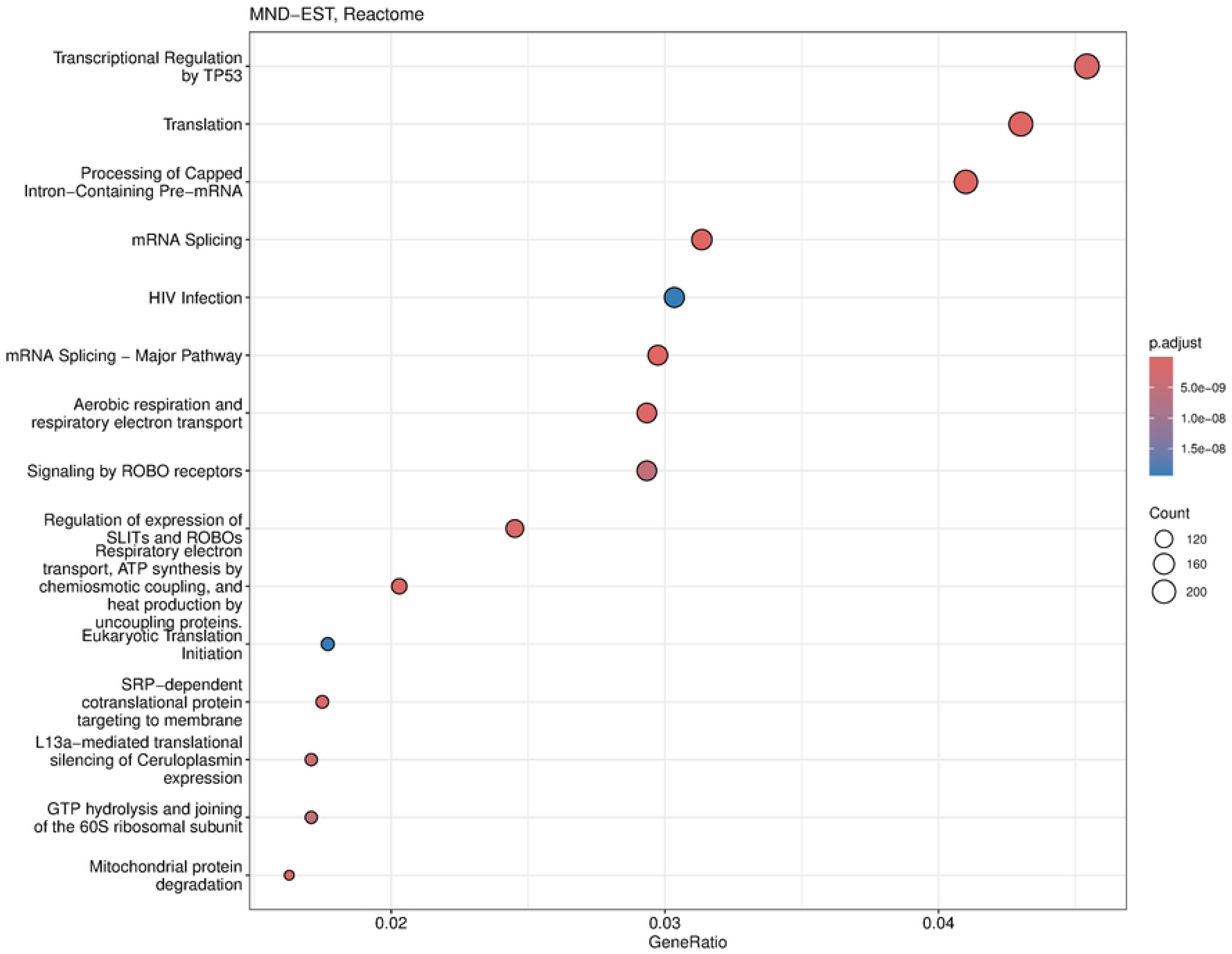
Dotplot of Reactome analysis based on the fold-change expression differences in the blood of MND patients. Top 15 the most significantly upregulated pathways are shown.

**Figure 7.**
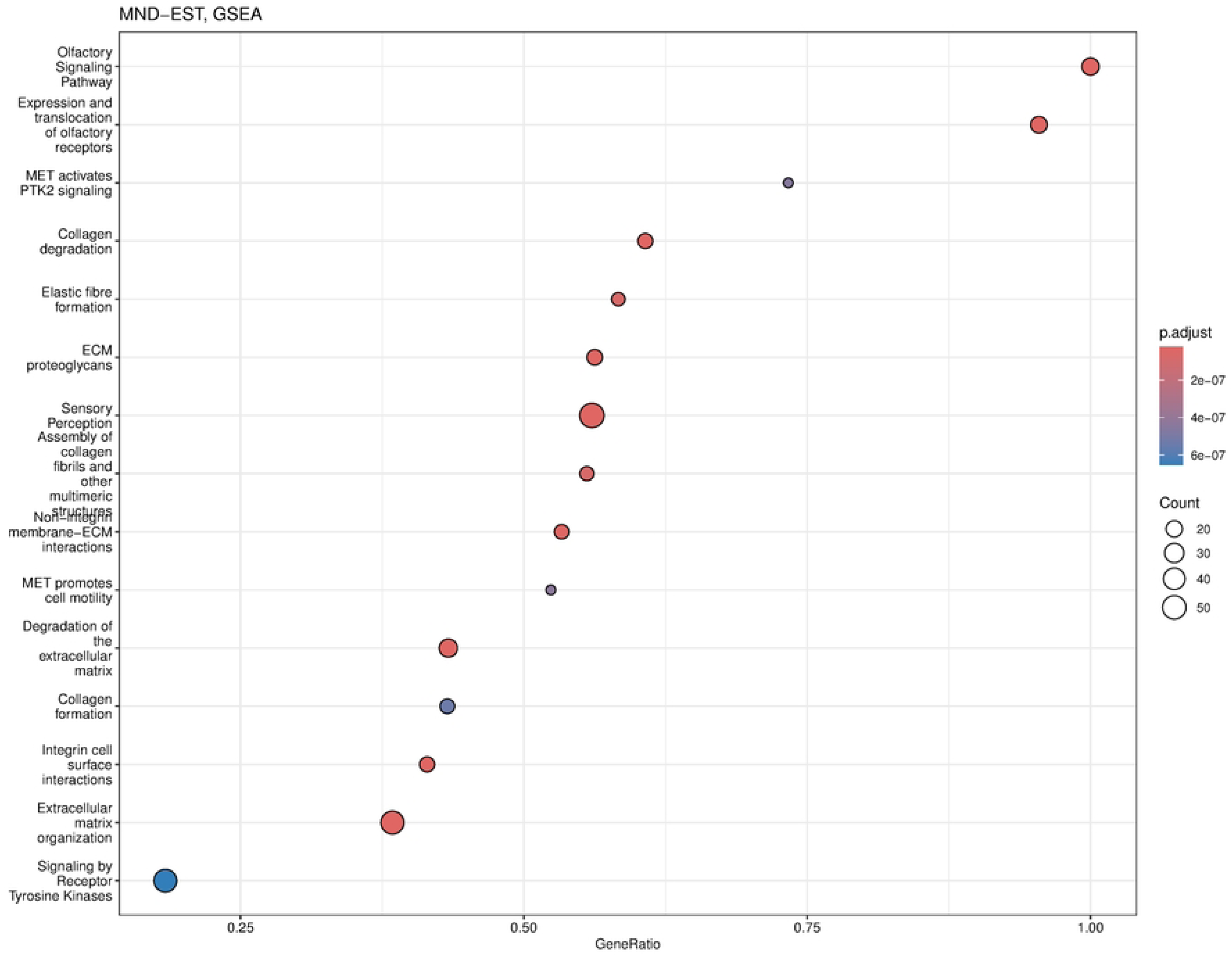
Dotplot of GSEA analysis based on the fold-change expression differences in the blood of MND patients. Top 15 the most significantly upregulated pathways are shown.

In summary, KEGG pathway analysis found statistically significant activation of the ALS pathway together with other neurodegeneration pathways. The findings from Reactome and GSEA added more details to the KEGG finding and identified several cellular pathways that can give a mechanistic understanding of the pathogenesis of MND.

## 4. Discussion

The current study presented a whole transcriptome analysis of the whole blood RNA from MND patients compared to age and sex-matched healthy controls. As a main finding, we identified 12,972 genes differentially expressed; 8,008 were upregulated, and 4,964 were downregulated in the blood of MND patients. Most remarkably, the heatmap based on these 12,972 genes was highly specific and separated MND from healthy controls. Therefore, we can conclude that the identified differentially expressed genes are specific for the MND status. This doesn’t mean that all of these genes are directly related to the pathogenesis of MND but instead reflects the complexity of the disease, where pathogenic changes are mixed with compensatory changes. However, this still shows that MND, while a CNS-specific disease, has remarkable changes in the blood transcriptomics, and blood could be a perfect source for the diagnostic biomarkers for MND.

Not all MND-specific genes were differentially expressed. C9orf72 is a gene with the highest genetic impact in MND, but it was not differentially expressed. C9orf72 is highly expressed in the blood with an average normalised count of 1,500. Therefore, the low expression level cannot explain the lack of significant differences. A similar observation is true for the SQSTM1, TARDBP, OPTN and PSEN1, all genes with high expression in the blood, but no difference in expression between MND and controls (Supplementary Figure 2). It is hard to understand why these genes did not show differential expression, but these genes have a mutation-specific effect, and in our cohort, we may not have mutations in these genes. This might be unlikely, as we have identified pathogenic repeat polymorphism for C9orf72 in one patient who has 1,000 repeats with a length of over 6,000 bp.

We saw significant differences in many MND-related genes. For instance, SOD1 was downregulated in MND patients. Similarly, ANG and ACSL5 were significantly downregulated in MND patients compared to controls. It is somewhat surprising that SOD1 is downregulated in MND patients as it is also assumed to form aggregates in sporadic patients [25–27]. At the same time, we couldn’t find a significant difference for the OPTN gene, another gene that has clear implications in MND pathology and had a very high expression level in blood. It is remarkable that while its aggregates are common for familial and sporadic MND forms, we could not detect significant differences in the expression of OPTN [28].

Our study is certainly not the first to analyse MND patients’ transcriptomes. One study analysed gene chips from whole blood RNA, finding 2,943 genes differentially expressed [17]. These authors did not find SOD1, C9orf72, SQSTM1, TARDBP, or OPTN to be differentially expressed; this study got similar results to ours. Other published studies have used selected cell fractions, like PBMCs or lymphoblastoid cells [16, 29–31]. The cell fractionation studies identified a much smaller number of differentially expressed genes, and their results are difficult to compare to our results as the approaches are quite different. However, one recent study used a machine learning approach to compare brain and blood transcriptomic data and identified three distinct clusters of the MND subtypes with potentially different pathological mechanisms [6]. These three pathogenic subtypes didn’t describe any particular MND mutation but rather the biological pathways that involved particular differentially expressed genes. The present study is based on blood transcriptome, and we have identified similar differentially expressed genes. While we couldn’t identify three distinctive subtypes, the heatmap of the 12,972 differentially expressed genes separated MND patients from controls. Moreover, for MND patients, we saw at least two clusters with specific gene expression profiles. Therefore, our study results seem to match the results of the study by Marriott et al [6]. The main finding is that gene expression profiles and RNA analysis could be used as a source for biomarkers and can have clinical utility in differentiating patients with distinctive pathogenetic mechanisms.

We identified that the most up-regulated gene, with logFC 23, in MND blood is the APOBEC3DE gene (Volcano plot in Figure 2). APOBEC3DE is located at 22q13.1 and is a cytidine deaminase gene family member. This gene is one of the APOBEC cluster family on chromosome 22 [32, 33]. APOBEC proteins are part of innate immunity, and they inhibit retroviruses by deaminating cytosine residues in retroviral cDNA [34]. Interestingly, APOBEC3DE also inhibits retrotransposition of the long interspersed element-1 (LINE-1) by interacting with ORF1p, a protein encoded by LINE-1 [35]. LINE-1 has been implicated in the pathogenesis of MND, and therefore, APOBEC3DE finding seems very relevant as they suppress LINE1 activity [36]. In addition, APOBEC proteins can induce somatic mutations into genomic DNA and promote the development of different diseases [37]. APOBEC proteins are also involved in the clearance of foreign DNA from human cells, implicating their role in the cellular defence system against mutations that make them very plausible in connection with the MND [38, 39]. Loss of the nuclear TDP-43 due to the cytoplasmic aggregation of the TDP-43 is associated with decondensation of the chromatin around LINE1 elements and increased activation or LINE1 with their retrotransposition. Upregulation of the APOBEC3DE might be an endogenous defence mechanism as it is a part of the innate response to retroviral activation [40].

Many differentially expressed genes are involved in splicing and RNA processing: RNU5A-1, RNU1-1, RNY3, and RNY1, to name some. Interestingly, these RNA synthesis and splicing-related genes are all upregulated in MND samples and not expressed in the blood of control samples at all. These are genes that have a high potential to become a blood biomarker for MND or help to predict the progression of the disease. While it is not clear how these genes participate in the pathogenesis of MND, splicing mutations and genes participating in splicing involvement in MND have been shown in many previous studies [41–43]. The results from blood transcriptomics were very uniform and showed the upregulation of several genes related to RNA synthesis and splicing, as also indicated in Figure 6.

The function of downregulated genes is more diverse, with possible common denominators being the extracellular matrix (ECM) organisation and remodelling (Figure 7). Reduced expression of CCDC80, COL1A1, COL1A2, MMP2, and TNFRSF11B indicates the ECM reorganisation also found in GSEA enrichment analysis (Figure 2). The expression of these genes was very low in MND samples and very high in the blood of controls, showing a highly significant logFC for these genes. Similarly, IGFBP5 almost lacked expression in the MND group and had very high expression in the blood of control subjects. Overexpression of the IGFBP5 in mice has induced axonopathy and sensory deficits similar to those seen in diabetic neuropathy [44]. The motor axon degeneration in these mice resembled the pathology seen in MND [44]. IGFBP5 has been shown to promote neuronal apoptosis in the experimental models and also in patients with spinal muscular atrophy and ALS [45–47]. When discussing these results, we have to consider the effect of MND itself on gene expression and not only the effect of genes on the disease. Most likely, the genes that are significantly downregulated and have very low expression levels in MND patients are the genes that are affected by the MND condition. The cluster of ECM organisation genes indicates the degeneration of the neurones and are the genes directly impacted by the MND. Stanniocalcin 2 (STC2) and thrombospondin 2 (THBS2) are genes that are related to organogenesis and tissue differentiation [48–50]. Interestingly, the proposed function of these genes is related to collagen genes and MMPs. Therefore, it seems that MND affects tissue reorganisation, and the genes that are required for tissue plasticity are downregulated. We can speculate that genes are not causative for the disease but are affected by the chronic disease condition and lead to enhanced degeneration of neurones.

## 5. Conclusions

We performed whole transcriptome analysis from the whole blood RNA and identified 12,972 genes differentially expressed between MND patients and controls. These gene expression changes have the potential to be used as biomarkers to diagnose MND and possibly to evaluate the progression of the disease and drug responsiveness in clinical trials. RNA-based biomarkers have excellent potential as they are quickly responding biomarkers and can be analysed by standardised methods. In conclusion, we were able to identify the characteristic blood gene expression profile of MND patients.

## Supplementary Materials

The following supporting information can be downloaded at: www.mdpi.com/xxx/s1, Figure S1: HeatmapMNDEST12972genesBloodRNAseq; Figure S2: AllBoxplots; Table S1: DescriptionofTheCohort; Table S2: Differentially expressed genes; Table S3: CompleteListofMNDGENES; Table S4: FC_MNDEST_KEGG; Table S5: FC_MNDEST_Reactome; Table S6: FC_MNDEST_GSEA.

## Author Contributions

Conceptualization, S.K. and P.T.; methodology, S.K., K.R, M.M, J.P.; software, S.K.; validation, S.K, A.L.P, Y.Y. and Z.Z.; formal analysis, S.K..; investigation, K.R, M.M, J.P.; resources, S.K. and P.T.; data curation, S.K.; writing—original draft preparation, S.K.; writing—review and editing, S.K., K.R, M.M, J.P., A.L.P, P.T.; visualization, S.K.; supervision, S.K.; project administration, S.K.; funding acquisition, S.K. and P.T. All authors have read and agreed to the published version of the manuscript.

## Institutional Review Board Statement

The study was conducted in accordance with the Declaration of Helsinki and approved by the Institutional Review Board (or Ethics Committee) of the University of Tartu 327/T-17 on 19.10.2019. the title of the approved study is “Amyotrophic lateral sclerosis (ALS) in Estonia: longitudinal epidemiological and molecular study”.

## Informed Consent Statement

Informed consent was obtained from all subjects involved in the study.

## Data Availability Statement

Data are deposited in the Gene Expression Omnibus under accession number GSE277709.

## Acknowledgments

The technical support from the Genomics Core Facility of Mudoch University is appreciated.

